# EpigeneticAgePipeline: an R package for comprehensive assessment of epigenetic age metrics from methylation microarrays

**DOI:** 10.1101/2024.10.24.620090

**Authors:** Stanislav Rayevskiy, Quinn Le, Julia Nguyen, Steven Q. Chen, Christina A. Castellani

## Abstract

Epigenetic age is a biological age estimate based on nuclear DNA methylation patterns. Epigenetic clocks measure biological age by analyzing predictable changes in DNA methylation sites associated with aging. This study introduces EpigeneticAgePipeline, an R package that streamlines the estimation of epigenetic age metrics including Horvath, Horvath skin and blood, Hannum, PhenoAge/Levine, GrimAge (V1, and V2), and DunedinPACE plus additional acceleration metrics based on all other clocks. Quality control includes detection p-value filtering (sample- and probe-level), bead-count thresholds, and Illumina quality control intensity checks. EpigeneticAgePipeline supports Illumina Infinium methylation microarrays (HumanMethylation27, HumanMethylation450, HumanMethylationEPIC/EPICv2, and Human Methylation Screening Array). It offers functionalities including data preprocessing, normalization, cell count imputation, residual generation accounting for principal components and batch effects, and extensive visualizations for improved interpretability. Validation was performed using GEO dataset GSE237561, confirming the accuracy of the pipeline. EpigeneticAgePipeline provides an integrated workflow from raw data to advanced statistical analyses and visualizations, improving usability over existing tools. In addition to the traditional clocks mentioned, the package also integrates a set of additional epigenetic age clocks (PedBE, Wu, TL, BLUP, and EN). Future updates will include emerging epigenetic age measures to maintain relevance in this evolving field.

## Background

The field of epigenomics has allowed for the understanding that environmental factors can significantly influence our genome and how particular regions of our genome are regulated and expressed. As the field has expanded, research into the association of epigenetic features, human health, disease risk, as well as aging has emerged, leading to the concept of epigenetic age and the creation of epigenetic age clocks. Epigenetic clocks are computational tools that estimate biological age by analyzing specific DNA methylation patterns that are associated with cellular aging across the genome. These clocks use measures of DNA methylation, chemical modifications that regulate gene expression without altering the underlying DNA sequence, to provide a look into the aging process. This is important because epigenetic age may be a better measure of an individual’s health status than chronological age alone. *EpigeneticAgePipeline* aims to build off the functionality of other well established R packages such as dnaMethyAge [1], methylclock [2], and minfi [3] to provide additional functionality and significantly improve usability.

The aim of this manuscript is to showcase the complete workflow performed by the package, from data preprocessing to visualization. This comprehensive approach emphasizes the seamless integration of functionalities, highlighting the distinct advantages of EpigeneticAgePipeline compared to other tools that typically require multiple separate steps to achieve similar outcomes.

What makes *EpigeneticAgePipeline* unique is that it integrates the entire workflow, including data pre-processing, epigenetic age calculations, and downstream visualization tasks/final analysis into one seamless pipeline. The package allows for direct input of raw intensity data (IDAT) files (27K, 450K, EPICv1/v2 and MSA microarray), which are then processed via sample and probe-level detection p-value filtering, bead-count thresholds at both the sample and probe levels, Illumina control intensity checks, and the removal of cross-reactive probes and sex probes [4–8]. Alternatively, users can input their own preprocessed methylation beta values. The package also allows for residual generation to account for cell count data and batch effects that arise during methylation data collection. Residual generation also includes the option to regress out principal components generated from the methylation dataset to account for additional sources of variation in the data. Cell proportion estimation is also included, allowing the user to specify tissue type and methodology.

Furthermore, *EpigeneticAgePipeline* provides a range of visualizations, including a variety of correlation and distribution plots that enhance the data’s interpretability. These features, combined with the foundational functionalities of other well-established R packages, allow *EpigeneticAgePipeline* to fill a unique niche in computational epigenetics.

### Implementation

*EpigeneticAgePipeline* generates five common epigenetic age clocks, which include Horvath clock [9], Horvath skin and blood clock [10], Hannum clock [11], PhenoAge/Levine clock [12] and GrimAge (V1 and V2 based on their published surrogate panels) [13,14]. We’ve also included the PC versions for each of these clocks [1]. In addition to these 5 common clocks, the pipeline also generates the PedBE clock [15], Wu’s clock [16], Telomere Length’s (TL) clock [17], BLUP clock [18], and EN clock [18]. Each one analyzes the methylation pattern of a unique set of CpG (Cytosine-phosphate-Guanine) sites, which allows for more specific estimations depending on the tissue type and/or phenotype. Along with epigenetic age estimation, the package allows for the generation of DunedinPACE [19], which reflects the pace of biological aging, and age acceleration for each of the generated clocks.

When raw IDATs are provided, the pipeline performs detection p-value filtering at two levels: (i) sample-level retention if the mean detection p-value is < 0.05, and (ii) probe-level retention only if the detection p-value is < 0.01 across passing samples. Bead-count filters exclude samples with > 0.05 low-bead observations (bead count < 3) and drop probes with > 0.05 low-bead observations among retained samples. Default parameters concerning detection p and bead count filtering can also be modified by the user. Illumina control intensity metrics are assessed via minfi, where samples with mean log2 intensity of mMed and uMed < 10.5 are removed [3]. The pipeline then removes known cross-reactive probes and sex-chromosome probes. For each of the microarray formats accepted (27K, 450K, EPICv1/v2, MSA), a separate list of previously identified cross reactive probes is removed during this process [20–23]. Currently there has been no formal study which compiled a list of cross-reactive probes for the MSA platform, however due to its similarity to EPICv2, we used EPICv2’s list.

To handle replicate probes in newer EPICv2/MSA array formats targeting the same legacy CpG, beta values are collapsed by probe-ID prefix using betasCollapseToPfx from the sesame R package, yielding a single averaged beta per CpG prior to clock computation [24].

To calculate cell proportion estimates, the package supports Constrained Projection (CP) and Robust Partial Correlations (RPC) style generation for adult blood, cord blood, saliva, placenta (first and third trimester), and frontal cortex cells [25–32]. Buccal cell proportions are also supported using the HEpiDISH (from EpiDISH R Package) hierarchical approach by first deconvolving epithelial, fibroblast and immune fractions with a solid-tissue reference, then partitioning the immune fraction into blood cell subtypes [26,33,34].

For all clocks except DunedinPACE, the package provides three variations of epigenetic age acceleration: (i) the difference between Epigenetic Age and chronological age, (ii) residuals from Epigenetic Age ~ Age + estimated cell proportions, and (iii) residuals from Epigenetic Age ~ Age. These are analogous to the age accelerations provided by methylclock [2].

The original Horvath clock was one of the first epigenetic clocks created and remains one of the “gold standard” epigenetic age measure. The clock uses a set of 353 CpG sites trained on 27K/450K arrays and is designed to be used on a wide range of tissue types. The second original epigenetic clock is the Hannum clock. This clock relies on a set of 71 CpG sites trained on 450K array and is primarily designed for use on blood samples. Both the Horvath and Hannum clocks have been shown to be significant predictors of age- and age-related conditions. After the creation of the Horvath clock, the Horvath skin and blood clock was developed using a set of 391 CpG sites trained on 450K/EPICv1 array types for better estimations from skin, blood, saliva, human fibroblasts, keratinocytes, buccal cell, endothelial cell and lymphoblastoid cell inputs. Another epigenetic age clock included is the Levine/PhenoAge clock which was created as a potential outperformer of other clocks concerning predictions of age-based phenotypic outcomes, it uses a set of 513 CpG sites trained on 27K/450K/EPICv1. GrimAge is a measure primarily designed for blood samples, with version 1 and 2 based on a set of 1030 CpG, difference being that version 2 uses an additional 2 DNAm (DNA methylation) surrogates. The surrogates are DNAm log(C-reactive protein) and DNAm log(Hemoglobin A1C), which allows GrimAge2 to outperform the first in mortality prediction [14]. GrimAge has been known to predict a high Mortality Hazard Ratio compared to other common epigenetic age measures. The PedBE clock uses 94 CpGs and was trained primarily on buccal epithelial samples (children and adolescents) profiled on 450K/EPIC platforms. It is optimized for pediatric tissues and has been used to track developmental timing and age acceleration in childhood cohorts. Wu’s clock uses 111 CpGs and was trained on pediatric blood using 27K and 450K arrays. It provides highly accurate age estimates in children and has been applied to investigate developmental and clinical outcomes in pediatric populations. The TL clock uses 140 CpGs trained on blood using 450K and EPIC arrays to estimate leukocyte telomere length. It has been used to identify associations between shorter telomere length and advancing age and relates to aging-related morbidity. The BLUP clock uses 319,607 CpGs trained on blood and saliva measured on 450K and EPIC arrays. It leverages genome-wide features to yield high-precision age prediction and robust performance across cohorts and tissues. The EN clock uses 514 CpGs trained on the same 450K/EPIC blood and saliva datasets as BLUP. The elastic-net (EN) clock provides a sparse, portable predictor with strong cross-platform accuracy and reliable performance in external data.

The DunedinPACE measure is a measure of the pace of biological aging, which has been known to show strong associations with phenotypic outcomes like morbidity, disability, and mortality. DunedinPace was developed for use in blood samples and uses a set of 173 CpG sites.

*EpigeneticAgePipeline* also incorporates residual generation to allow for correction of important confounding variables. Principal component analysis is performed on beta values, and residuals are generated adjusting for the first 5 PCs (default) generated on the methylation data. Samples that are beyond 3 times (default) the median absolute deviation in any of the first PCs are marked as outliers. Residual generation also incorporates adjustments for technical effects that commonly arise from methylation array measurements. The user may provide data for array variables such as Row, Column, Slide, and Batch. Depending on which are given, the model will dynamically construct a suitable formula for the regression (see Figure 1). There is also a built-in option to compute and adjust for cell count proportion estimates. Residuals generated when chronological age is set as a predictor for biological age effectively gives a measure of age acceleration.

**Figure 1:**
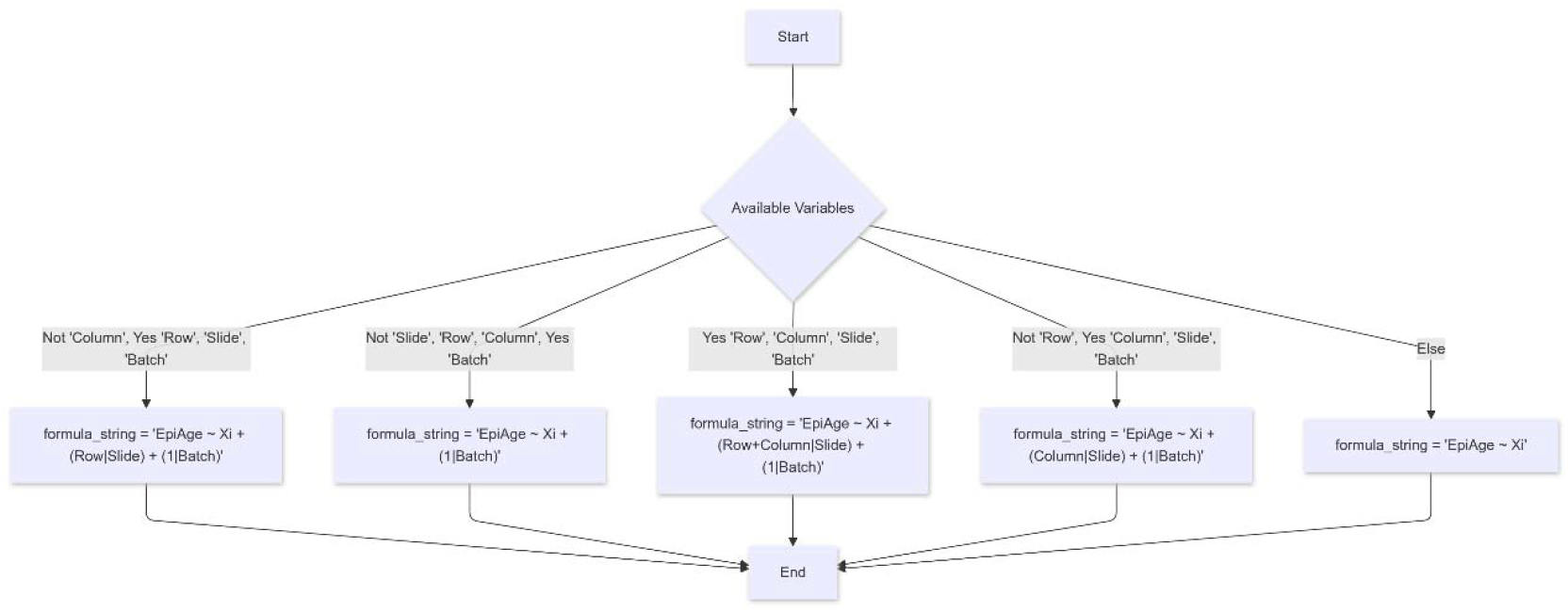
Flowchart illustrating formula construction for the linear model during residual generation (*generateResiduals* function). *Xi* represents all other user provided covariates. This schematic is based off presence of covariates “Row”, “Column”, “Slide”, and “Batch”.

A series of plots and visualizations are also included in the final output to present the correlations between epigenetic age measures and chronological age as well as the distribution of epigenetic age estimates for every sample. Raw data is exported as.txt file which can be used for further analysis.

## Results

The package’s *main* function is the primary entry point for epigenetic age analysis. Users provide the directory containing methylation and covariate data, if available (e.g., age, sex, smoking status, batch effects, etc.), along with options for normalizing beta values, selecting the methylation array type, and processing raw or precomputed data. The function performs preprocessing steps, cell count generation, and the creation of a file containing beta values for downstream analysis. This function also supports parallel processing for efficient handling of large datasets and allows users to specify whether to use adult or cord blood assumptions during cell count generation.

The *generateResiduals* function is designed for advanced analysis following epigenetic age estimation. This function builds a linear model based on user-specified variables or principal components (PCs) and outputs the corresponding residuals, which can be used for further investigation of epigenetic age-related patterns. Users can input custom formulas for the model, specify the type of methylation array used, and include additional covariates such as cell counts in the analysis. The function also supports the detection and removal of highly correlated variables (if enabled) and identifies outliers using median absolute deviation (MAD). As with the main function, this process can leverage parallel processing for efficiency and accommodates both adult and cord blood sample assumptions during cell count generation. To view a complete outline of the workflow for either the main or generateResiduals function, see Figure 2/3.

**Figure 2:**
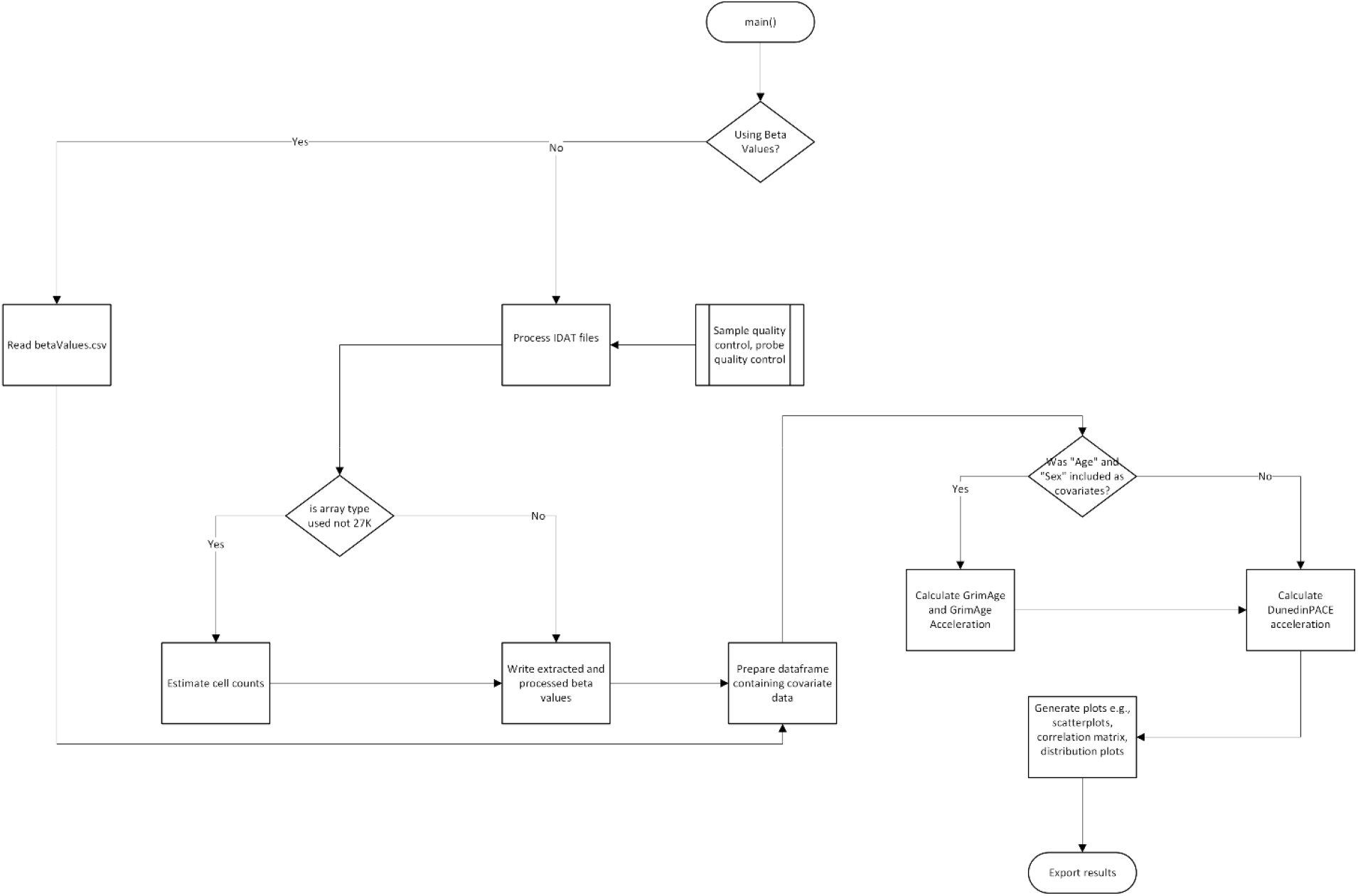
Workflow diagram illustrating the primary functionalities of the main() function. Flowchart for the main() function, which outlines the step-by-step process from data input to epigenetic age estimation and visualization, highlighting options for IDAT or beta value input and downstream processing.

**Figure 3:**
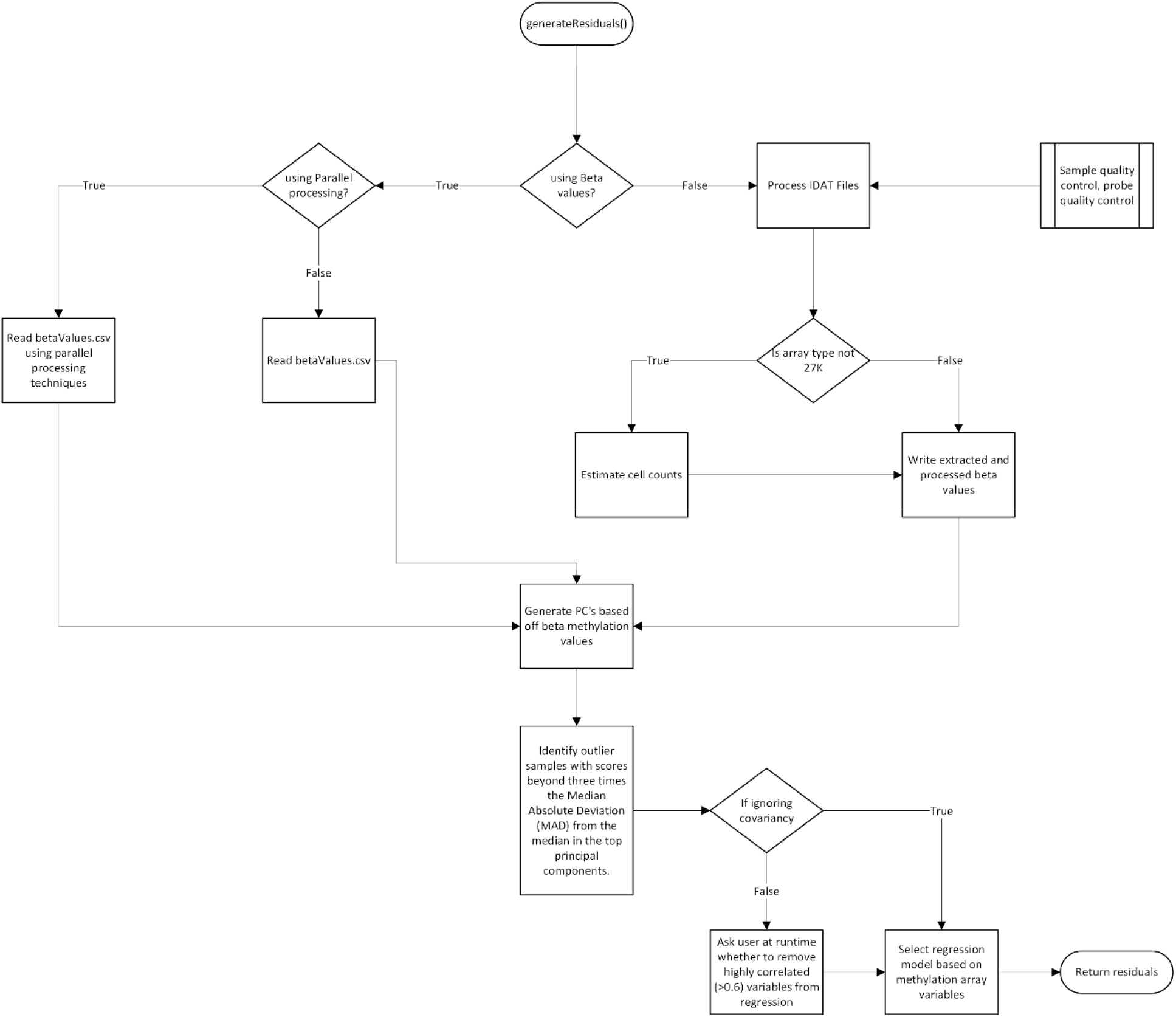
Workflow diagram illustrating the primary functionalities of the generateResiduals() function. Flowchart for the generateResiduals() function, showcasing the process for residual generation, including data preprocessing, principal component analysis, outlier detection, and regression model construction.

To ensure the robustness of our pipeline, we first validated its implementation using the publicly available Gene Expression Omnibus (GEO) dataset GSE237561 [35]. This dataset contains nuclear DNA methylation data for 126 samples—92 measured using the 450K array and 34 using the EPIC platform. Since both methylation and phenotypic data are publicly available, we were able to compare our results against previously published findings. Specifically, a prior study investigating the relationship between clozapine use and epigenetic changes utilized this dataset to calculate PhenoAge/Levine and DunedinPACE epigenetic age measures [36]. These epigenetic markers were incorporated into linear mixed models to assess whether clozapine use predicts epigenetic age or age acceleration. To validate our pipeline, we used all 126 samples and compared our computed PhenoAge/Levine and DunedinPACE measures against the published results. Our package generates these estimates using the methylclock and dnaMethyAge packages, respectively, this comparison therefore seeks to confirm the accuracy of our implementation. The full validation results can be found in the *Supplementary Information*. Following this validation, we applied our pipeline specifically to the 34 EPIC-measured samples for visualization. The covariates included in this analysis were sex, age, smoking score and number of days on clozapine. Figure 4, as well as all *Supplementary Figures* (except validation) were generated using our EpigeneticAgePipeline on the EPIC-measured samples from the GSE237561 dataset.

**Figure 4:**
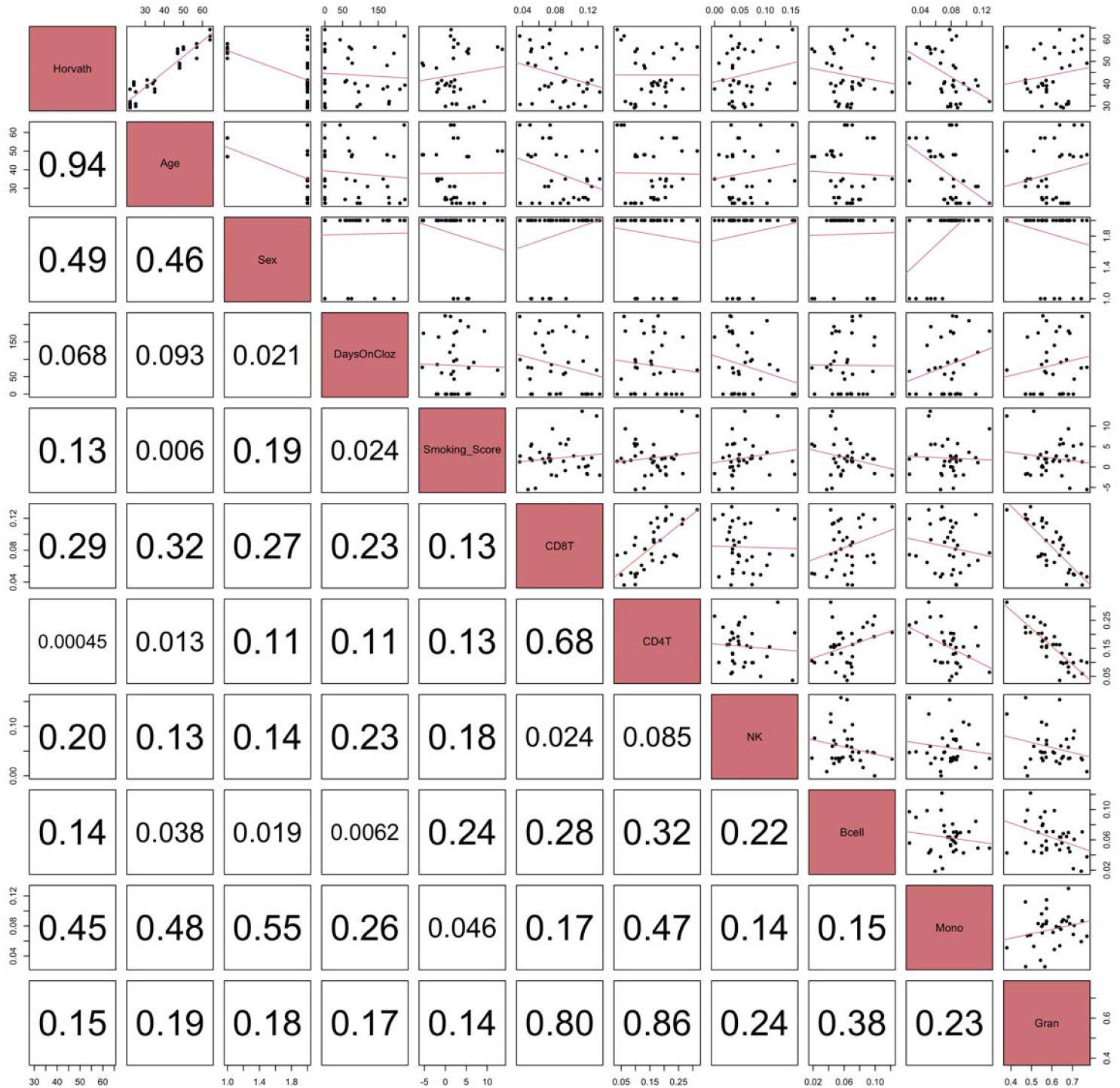
A correlation matrix generated for Horvath epigenetic age from the GSE237561 dataset. The boxes below the diagonal showcase the magnitude of the correlation between each respective pair of variables. The boxes above the diagonal showcase a scatterplot of pairwise variables with a linear trendline, giving the directionality and strength of the correlation. The *main* function will output an analogous plot for each measure of epigenetic age if covariates have been included.

### Inputs

The package accepts methylation data either in the form of a.csv file containing methylation beta values for each respective sample, or a directory containing raw IDAT files.

### Outputs

The package outputs raw epigenetic age/acceleration values (.txt and.md, Table S1), raw residual results (.csv), age and epigenetic age correlation plots (Figure S2), matrix correlation plots (.png, Figure 3), and an epigenetic age distribution plot (.png, Figure 3). A clock-coverage summary table is also outputted, reporting for each clock/surrogate the total CpGs, missing CpGs, and percent missing (Table S2). Examples of all the plots can be found in the *Supplementary Information* files.

Specific information on installation procedures and file specificities, as well as vignettes are located within the GitHub readme [37].

## Conclusions

*EpigeneticAgePipeline* provides a comprehensive and user-friendly method to generate epigenetic age measures for those studying the effects of biological aging. Our pipeline better accounts for potential batch effects and confounding variables to minimize the effect of potential biases on the results in studies concerning biological aging. Our pipeline provides a complete workflow for epigenetic age estimation, starting with raw methylation data all the way to the generation of epigenetic age measures, incorporating a variety of quality control methods and visualizations.

*EpigeneticAgePipeline* serves as a valuable tool for investigating the relationships between epigenetic age measures and a wide range of health outcomes. Future updates will include the addition of emerging epigenetic aging measures/methods to ensure the pipeline remains relevant to this expanding field.

## Availability and Requirements

Project Name: EpigeneticAgePipeline

Project Homepage: https://github.com/CastellaniLab/EpigeneticAgePipeline

Operating Systems: Platform independent

Programing Language: R

Other Requirements: R 4.2+

License: CC0 1.0

Any restrictions to use by non-academics: No

## List of Abbreviations

GEO: Gene Expression Omnibus
IDAT: Intensity data file
PC: Principal Component
MAD: Median Absolute Deviation
CpG: Cytosine-phosphate-Guanine

## Declarations

### Ethics approval and consent to participate

Not applicable, this study used publicly available datasets and did not involve any human participants, human tissue, or human data collected by the authors.

### Consent for publication

Not applicable.

### Availability of data and materials

The datasets generated and/or analyzed during the current study are available in the Gene Expression Omnibus repository, accession code GSE237561, https://www.ncbi.nlm.nih.gov/geo/query/acc.cgi?acc=GSE237561

### Competing Interests

The authors declare that they have no competing interests

### Funding

This work was supported by start-up funds to CC from The Department of Pathology and Laboratory Medicine at Western University. It was further supported by institutional summer support (USRI) from Western University to SR.

### Authors’ contributions

SR led the software development portion of the project and contributed to testing and manuscript preparation. QL was involved in extensive software testing and co-wrote the manuscript. JN contributed to code development and provided critical feedback during the early stages of the project. SC assisted with testing and provided additional validation support. CC conceptualized and supervised the project, provided funding, and oversaw all aspects of development and manuscript preparation. All authors read and approved the final manuscript.

## Acknowledgements

We acknowledge the Digital Research Alliance and the Canadian Foundation for Innovation J. Evans Leaders Fund grant #43481 for the support of the computing infrastructure used throughout our analyses.

## Notes

https://github.com/CastellaniLab/EpigeneticAgePipeline

### Competing Interest Statement

The authors have declared no competing interest.

### Summary of Updates

We have added a number of new functions to the package

