## Supplementary Materials for "EpigeneticAgePipeline: an R package for comprehensive assessment of epigenetic age metrics from methylation microarrays"

### Supplementary Material

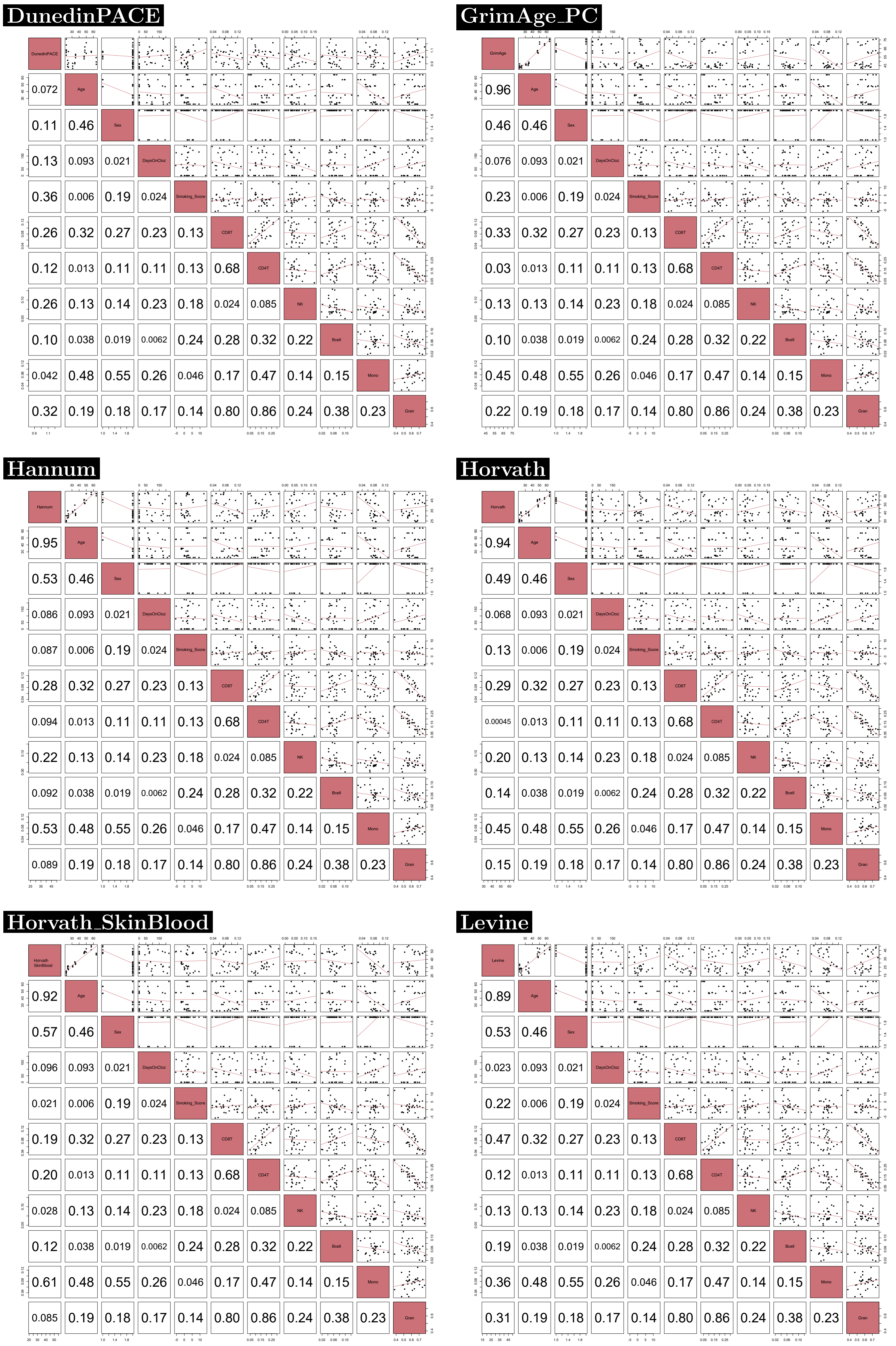


Figure 1: Example set of correlation matrices for each epigenetic age and epigenetic age acceleration measure with relevant variables from the GSE237561 dataset (using samples measured via EPIC). Each matrix is generated by EpigeneticAgePipeline as a separate .png file.

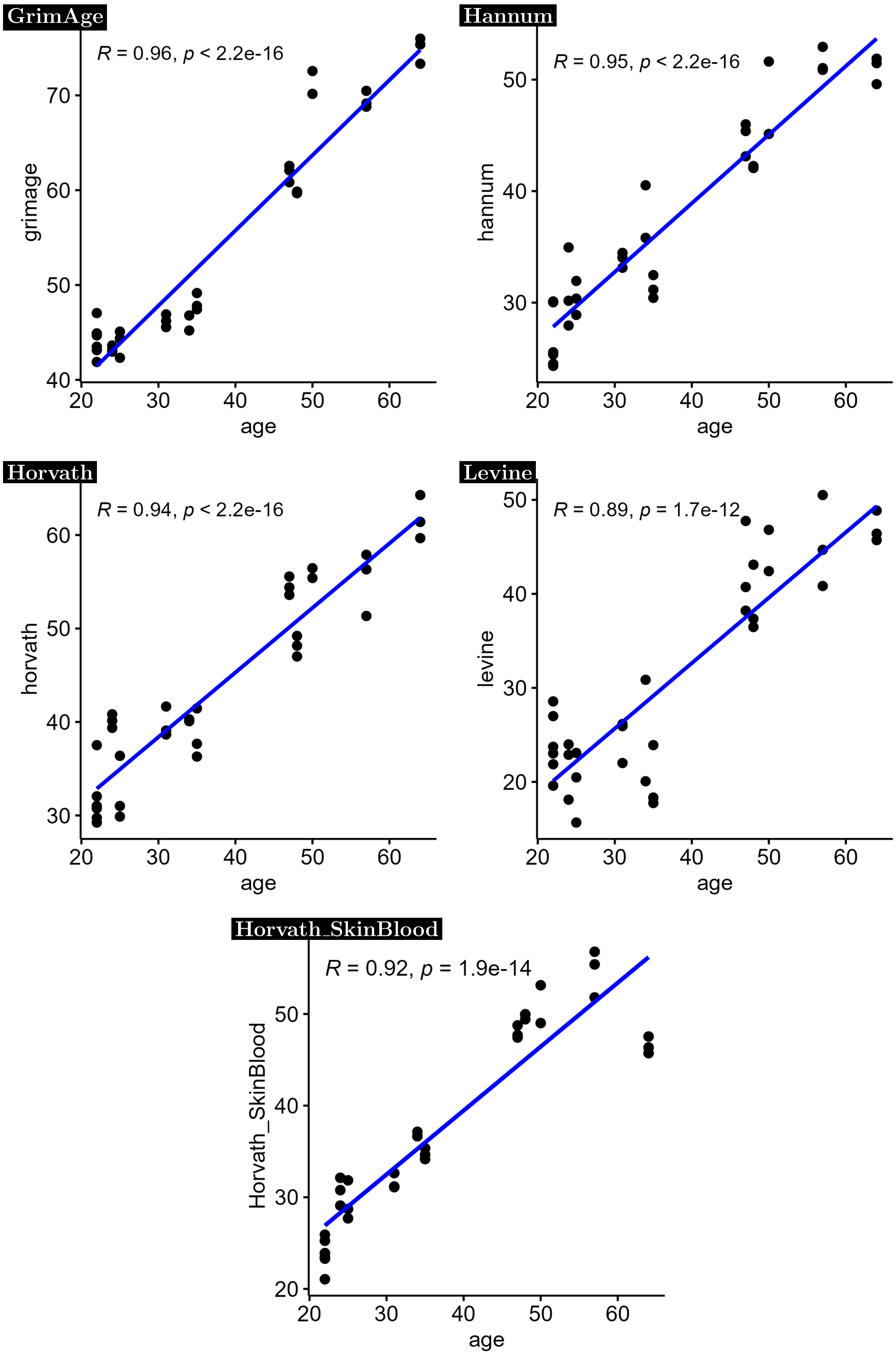


Figure 2: Example set of scatter plots with each epigenetic age measure plotted against chronological age. Each plot is generated separately by EpigeneticAgePipeline as a .png file. EPIC-measured samples from the GSE237561 dataset.

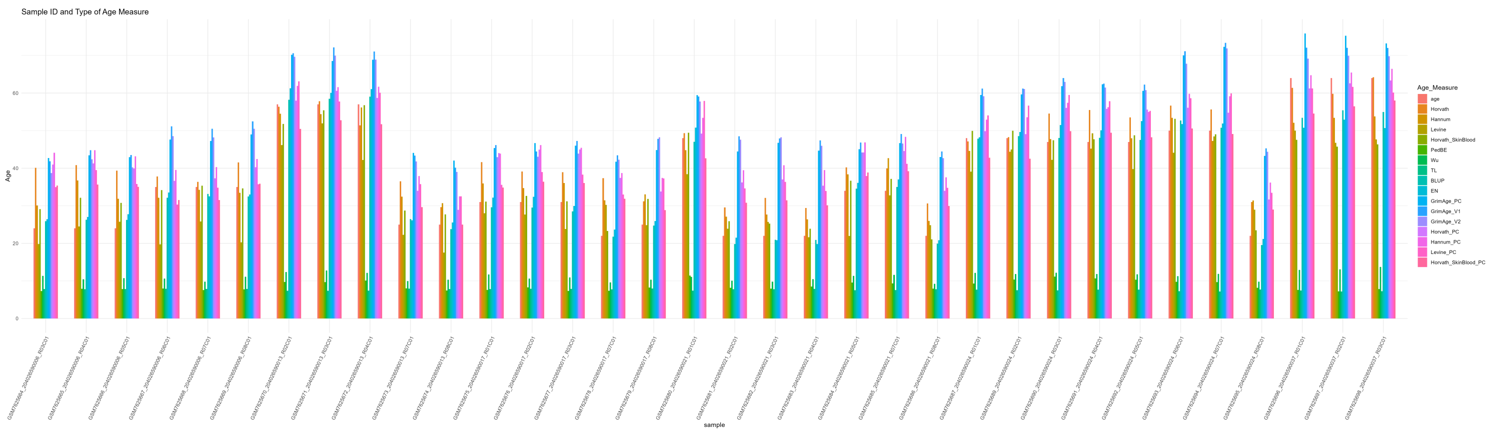

Figure 3: Example distribution plot which includes all possible measures of epigenetic age and chronological age. Exported as a .png file by EpigeneticAgePipeline. EPIC-measured samples from the GSE237561 dataset.

| id | Horvath | ageAcc.Horvath | ageAcc2.Horvath | Hannum | Horvath_SkinBlood | DunedinPACE | age |
|---|---|---|---|---|---|---|---|
| GSM7625664_204026590006_R03C01 | 40.09260526 | 16.09260526 | 5.844681576 | 30.08829652 | 29.09406948 | 0.855554556 | 24 |
| GSM7625665_204026590006_R04C01 | 40.82214834 | 16.82214834 | 6.574224659 | 36.75064772 | 32.12505905 | 0.886691227 | 24 |
| GSM7625666_204026590006_R05C01 | 39.3380305 | 15.3380305 | 5.090106818 | 31.8647584 | 30.78048325 | 0.94703597 | 24 |
| GSM7625667_204026590006_R06C01 | 37.7909955 | 2.790995495 | -4.074113658 | 32.14571792 | 34.16796129 | 1.026481493 | 35 |
| GSM7625668_204026590006_R07C01 | 36.37639297 | 1.376392967 | -5.488716186 | 34.24299389 | 35.352135 | 0.988684322 | 35 |
| GSM7625669_204026590006_R08C01 | 41.54186636 | 6.541866361 | -0.323242793 | 33.45145426 | 34.64390906 | 1.03950683 | 35 |
| GSM7625670_204026590013_R02C01 | 56.35177367 | -0.64822633 | -0.747706431 | 54.51798532 | 51.80183538 | 1.081416853 | 57 |
| GSM7625671_204026590013_R03C01 | 57.82392839 | 0.823928388 | 0.724448287 | 54.39371308 | 55.41039028 | 1.068924259 | 57 |
| GSM7625672_204026590013_R04C01 | 51.40533931 | -5.594660691 | -5.694140792 | 56.15040509 | 56.79220807 | 1.0772605 | 57 |
| GSM7625673_204026590013_R07C01 | 36.50230847 | 11.50230847 | 1.561913385 | 32.42548529 | 28.72552692 | 0.91278222 | 25 |
| GSM7625674_204026590013_R08C01 | 29.6487312 | 4.648731196 | -5.291663891 | 30.70292907 | 27.70010961 | 0.848163495 | 25 |
| GSM7625675_204026590017_R01C01 | 41.60606525 | 10.60606525 | 2.510841727 | 35.95724267 | 31.1066392 | 0.828741573 | 31 |
| GSM7625676_204026590017_R02C01 | 39.11027009 | 8.11027009 | 0.015046564 | 34.7211365 | 32.63816859 | 0.905680785 | 31 |
| GSM7625677_204026590017_R03C01 | 38.93379022 | 7.933790224 | -0.161433302 | 36.05497447 | 31.17273049 | 0.863070524 | 31 |
| GSM7625678_204026590017_R07C01 | 37.34721872 | 15.34721872 | 4.484237858 | 31.43739141 | 23.30073897 | 1.029797375 | 22 |
| GSM7625679_204026590017_R08C01 | 31.19800185 | 6.198001851 | -3.742393235 | 33.04506419 | 31.84846382 | 1.082512818 | 25 |
| GSM7625680_204026590021_R01C01 | 49.34258189 | 1.342581893 | -1.524655547 | 44.79570422 | 49.42421577 | 1.15197938 | 48 |
| GSM7625681_204026590021_R02C01 | 29.57199196 | 7.571991964 | -3.290988902 | 27.09422227 | 25.91437954 | 1.091249149 | 22 |
| GSM7625682_204026590021_R03C01 | 32.09795929 | 10.09795929 | -0.765021579 | 27.67065827 | 25.25819037 | 1.271073982 | 22 |
| GSM7625683_204026590021_R04C01 | 29.41165714 | 7.411657137 | -3.451323729 | 26.38776038 | 23.90729191 | 1.178856026 | 22 |
| GSM7625684_204026590021_R05C01 | 40.22042893 | 6.220428926 | -0.95220882 | 38.35513842 | 36.65659277 | 0.898388046 | 34 |
| GSM7625685_204026590021_R07C01 | 39.94881972 | 5.948819717 | -1.223818029 | 42.68278033 | 37.14246273 | 0.916559158 | 34 |
| GSM7625686_204026590021_R08C01 | 30.61530413 | 8.615304134 | -2.247676733 | 25.97858146 | 21.05502613 | 1.109341634 | 22 |
| GSM7625687_204026590024_R01C01 | 47.12700174 | -0.872998263 | -3.740235704 | 44.62126033 | 49.94020401 | 1.074309726 | 48 |
| GSM7625689_204026590024_R02C01 | 48.27267995 | 0.27267995 | -2.59455749 | 44.39054475 | 49.97322345 | 1.104276009 | 48 |
| GSM7625690_204026590024_R03C01 | 54.53674676 | 7.536746757 | 4.361980723 | 47.72835024 | 47.43278829 | 1.014802465 | 47 |
| GSM7625691_204026590024_R04C01 | 55.47904351 | 8.479043511 | 5.304277478 | 45.21076717 | 47.69100077 | 0.999165486 | 47 |
| GSM7625692_204026590024_R05C01 | 53.53381225 | 6.533812247 | 3.359046213 | 47.98272781 | 48.75344426 | 0.971099091 | 47 |
| GSM7625693_204026590024_R06C01 | 56.64984569 | 6.649845689 | 4.397665436 | 53.40175532 | 53.13307465 | 1.05346851 | 50 |
| GSM7625694_204026590024_R07C01 | 55.65307515 | 5.653075152 | 3.400894898 | 47.23181924 | 49.009247 | 1.210594457 | 50 |
| GSM7625695_204026590024_R08C01 | 30.99484761 | 8.994847613 | -1.868133253 | 31.43222898 | 23.486856 | 1.068861182 | 22 |
| GSM7625696_204026590037_R01C01 | 61.39938056 | -2.600619445 | -0.547399392 | 52.07367784 | 47.54001616 | 0.991021773 | 64 |
| GSM7625697_204026590037_R02C01 | 59.7794099 | -4.220590103 | -2.167370051 | 53.41986763 | 45.72042929 | 0.925105684 | 64 |
| GSM7625698_204026590037_R03C01 | 64.21421384 | 0.214213843 | 2.267433895 | 53.77038727 | 46.35112286 | 0.89079409 | 64 |

Table 1: Example output table including a subset measures of epigenetic age and epigenetic age acceleration for each sample, exported as .txt. EPIC-measured samples from the GSE237561 dataset.

| clock | Cpgs_in_clock | missing_CpGs | percentage |
|---|---|---|---|
| Horvath | 353 | 23 | 6.5 |
| Horvath_PC | 78464 | 2131 | 2.7 |
| Hannum | 71 | 9 | 12.7 |
| Hannum_PC | 78464 | 2131 | 2.7 |
| Levine | 513 | 4 | 0.8 |
| Levine_PC | 78464 | 2131 | 2.7 |
| Horvath_SkinBlood | 391 | 8 | 2 |
| Horvath_SkinBlood_PC | 78464 | 2131 | 2.7 |
| PedBE | 94 | 0 | 0 |
| Wu | 111 | 6 | 5.4 |
| TL | 140 | 23 | 16.4 |
| BLUP | 319607 | 9972 | 3.1 |
| EN | 514 | 10 | 1.9 |
| DunedinPACE | 173 | 16 | 9.2 |
| GrimAge_PC | 78464 | 2131 | 2.7 |
| DNAmADM_GrimAgeV1 | 186 | 46 | 24.7 |
| DNAmB2M_GrimAgeV1 | 91 | 12 | 13.2 |
| DNAmCystatinC_GrimAgeV1 | 87 | 15 | 17.2 |
| DNAmGDF15_GrimAgeV1 | 137 | 14 | 10.2 |
| DNAmLeptin_GrimAgeV1 | 187 | 55 | 29.4 |
| DNAmPACKYRS_GrimAgeV1 | 172 | 23 | 13.4 |
| DNAmPAI1_GrimAgeV1 | 211 | 40 | 19 |
| DNAmTIMP1_GrimAgeV1 | 42 | 6 | 14.3 |
| DNAmADM_GrimAgeV2 | 186 | 46 | 24.7 |
| DNAmB2M_GrimAgeV2 | 91 | 12 | 13.2 |
| DNAmCystatinC_GrimAgeV2 | 87 | 15 | 17.2 |
| DNAmGDF15_GrimAgeV2 | 137 | 14 | 10.2 |
| DNAmLeptin_GrimAgeV2 | 187 | 55 | 29.4 |
| DNAmlogA1C_GrimAgeV2 | 86 | 16 | 18.6 |
| DNAmlogCRP_GrimAgeV2 | 132 | 27 | 20.5 |
| DNAmPACKYRS_GrimAgeV2 | 172 | 23 | 13.4 |
| DNAmPAI1_GrimAgeV2 | 211 | 40 | 19 |
| DNAmTIMP1_GrimAgeV2 | 42 | 6 | 14.3 |

Table 2: Example output table showing clock coverage for a given input. EPIC-measured samples from the GSE237561 dataset.

| | Beta Value | SE Value | P Value |
|---|---|---|---|
| PhenoAge | 2.18 | 1.66 | 0.19 |
| PhenoAge Ref | 1.75 | 1.52 | 0.25 |
| DunedinPACE | 0.056 | 0.026 | 0.03 |
| DunedinPACE Ref | 0.042 | 0.025 | 0.09 |

Table 3: This analysis compares the results of mixed effects regression models predicting epigenetic age based on time spent on clozapine (in years), using both reference/dataset epigenetic measures and our package-generated epigenetic measures. The model includes all samples from the GSE237561 dataset and is defined as: PredictedAge $\sim$ timeOnClozapine + Age + Sex + (1 | Individual_ID) + (1 | Institute).

| | CD8T | CD4T | NK | Bcell | Mono | Gran |
|---|---|---|---|---|---|---|
| CD8T Ref | 1 | 0.44 | 0.01 | 0.4 | -0.12 | -0.66 |
| CD4T Ref | 0.44 | 1 | 0.11 | 0.19 | -0.36 | -0.73 |
| NK Ref | 0.02 | 0.1 | 1 | 0.22 | -0.04 | -0.43 |
| Bcell Ref | 0.39 | 0.19 | 0.22 | 1 | 0.07 | -0.61 |
| Mono Ref | -0.11 | -0.37 | -0.05 | 0.07 | 1 | -0.06 |
| Gran Ref | -0.66 | -0.73 | -0.43 | -0.06 | -0.06 | 1 |

Table 4: Correlations between package-generated cell proportions and reference/dataset data, assuming adult peripheral blood samples. Uses all samples from the GSE237561 dataset.
